# Entropic control of the free energy landscape of an archetypal biomolecular machine

**DOI:** 10.1101/2022.10.03.510626

**Authors:** Korak Kumar Ray, Colin D. Kinz-Thompson, Jingyi Fei, Bin Wang, Qiao Lin, Ruben L. Gonzalez

## Abstract

Biomolecular machines are complex macromolecular assemblies that utilize thermal and chemical energy to perform essential, multistep, cellular processes. Despite possessing different architectures and functions, an essential feature of the mechanisms-of-action of all such machines is that they require dynamic rearrangements of structural components. Surprisingly, biomolecular machines generally possess only a limited set of such motions, suggesting that these dynamics must be repurposed to drive different mechanistic steps. Although ligands that interact with these machines are known to drive such repurposing, the physical and structural mechanisms through which ligands achieve this remain unknown. Using temperature-dependent, single-molecule measurements analyzed with a time-resolution-enhancing algorithm, here we dissect the free energy landscape of an archetypal biomolecular machine, the bacterial ribosome, to reveal how its dynamics are repurposed to drive distinct steps during ribosome-catalyzed protein synthesis. Specifically, we show that the free energy landscape of the ribosome encompasses a network of allosterically coupled structural elements that coordinates the motions of these elements. Moreover, we reveal that ribosomal ligands which participate in disparate steps of the protein synthesis pathway repurpose this network by differentially modulating the structural flexibility of the ribosomal complex (i.e., the entropic component of the free energy landscape). We propose that such ligand-dependent entropic control of free energy landscapes has evolved as a general strategy through which ligands may regulate the functions of all biomolecular machines. Such entropic control is therefore an important driver in the evolution of naturally occurring biomolecular machines and a critical consideration for the design of synthetic molecular machines.

## INTRODUCTION

The structural motions of biomolecules are essential components of their function (*1–3*). While the contributions that such structural dynamics make to the reaction pathways of a number of small enzymes have been well studied, their role in the mechanisms of large biomolecular complexes remains underexplored. Such biomolecular ‘machines’ play essential roles in the cell, driving processes as fundamental as DNA replication, RNA transcription, messenger RNA (mRNA) splicing, and protein synthesis (*4*). A unique, defining feature of these machines is that they utilize nano-scale structural rearrangements to convert thermal and chemical energy into molecular-level mechanical work (*5,6*). The dynamics of these rearrangements are modulated by the interactions of the biomolecular machines with a host of different ligands, including substrates, inhibitors, and co-factors, in order to drive and regulate their functions (*7–11*). Unfortunately, technical challenges to studying the structural dynamics of large biomolecular complexes have thus far precluded an understanding of the physical and structural bases through which ligands exploit them to direct the functions of biomolecular machines.

Over the past twenty years, we have developed the capability to study the structural dynamics of an archetypal biomolecular machine, the bacterial ribosome, responsible for translating messenger RNAs (mRNAs) into proteins. Specifically, we have established a reconstituted *in vitro* translation system composed of ribosomes and other translation components purified from *Escherichia coli* (*E. coli*) (*12,13*). Using this system, we have developed numerous single-molecule fluorescence resonance energy transfer (smFRET) (*14–17*) signals that report on the conformational dynamics of the translating ribosome (*10,18–21*). Notably, two of these signals report on a fundamental, compound conformational change of the ribosome that is integral to many of the mechanistic steps of translation (*22*) (Fig. 1 and Fig. S1). Utilizing these and similar signals, we and others have shown that various ribosomal ligands can modulate the rate of this conformational change (*10,18,23–30*). In particular, transfer RNAs (tRNAs), the set of adaptor molecules that deliver amino acid substrates to the ribosome in the order specified by the nucleotide sequence of the mRNA, have been shown to modulate the dynamics of the ribosome. Remarkably, tRNAs differentially modulate ribosome dynamics in a way that depends on the identity, post-transcriptional modification status, and aminoacylation state of the tRNA (*10,18,23,25,30*). With this understanding, here we have used ribosomal complexes (RCs) that either lack or carry one of two classes of bound tRNAs as a model system. Combined with two novel technological advances (see below), this model system has provided us with a unique opportunity to investigate the physical and structural mechanisms through which the binding of different ligands (*i*.*e*., tRNAs) to a biomolecular machine (*i*.*e*., the bacterial ribosome) differentially modulate the conformational dynamics of the machine in order to direct its biological function (*i*.*e*., mRNA translation).

**Figure 1.**
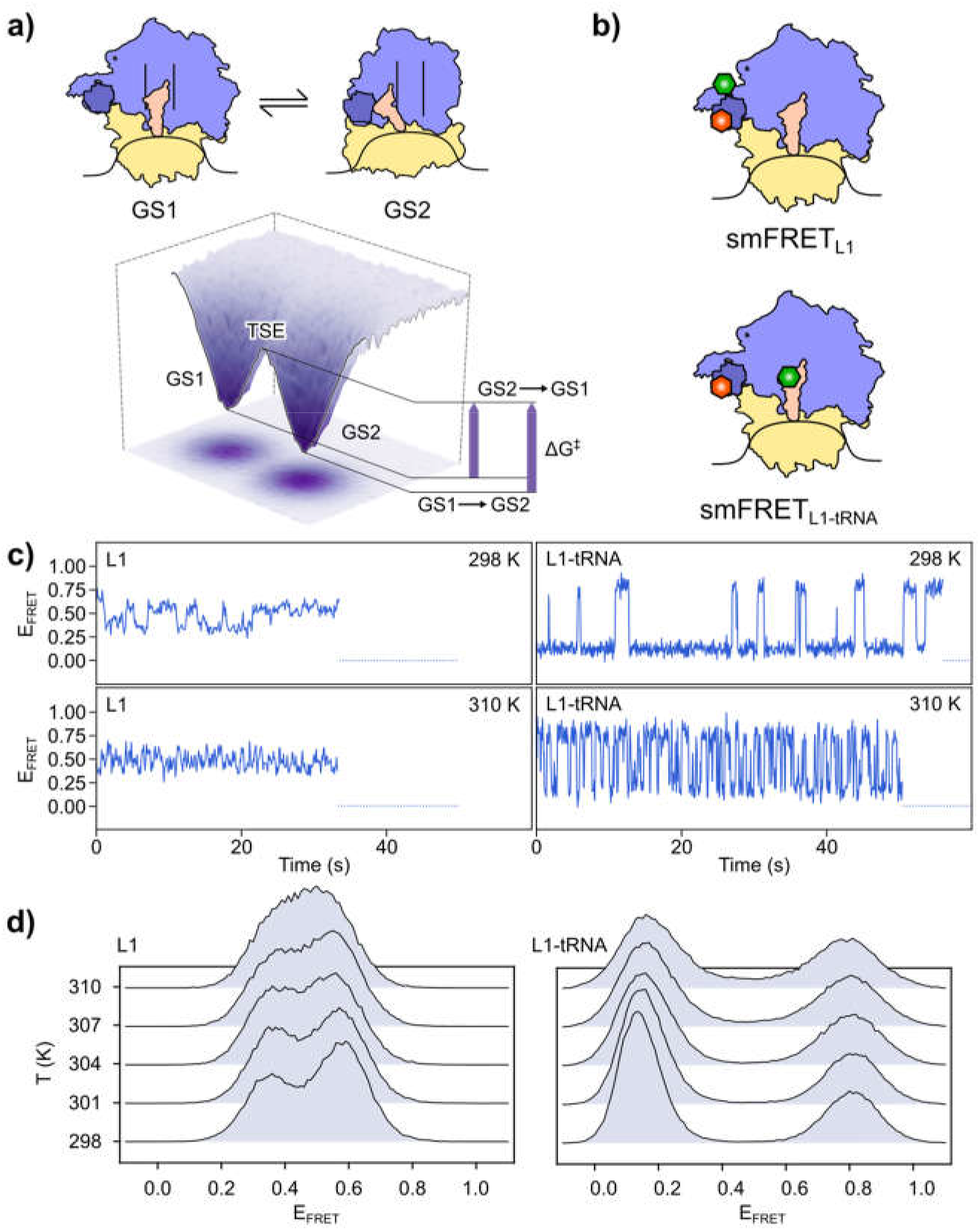
Temperature-dependent smFRET studies of the GS1⇌ GS2 equilibrium. **(a)** Structural cartoon representation of the GS1⇌ GS2 equilibrium in RC^Phe^ (above). The ribosomal small subunit is shown in beige, the ribosomal large subunit in purple, the L1 stalk in dark purple, the tRNA^Phe^ in orange, and the mRNA in black. The black lines in the large subunit demarcate the three different tRNA binding sites on the large subunit, two of which the tRNA moves between in transitions between GS1 and GS2. Representation of the free energy landscape (below) consisting of the wells (GS1 and GS2), barriers (TSE), and barrier heights (ΔG^‡^, labelled for the barriers traversed by the GS1→ GS2 and GS2→ GS1 transitions, respectively) that govern this equilibrium. **(b)** Positions of the donor (green) and acceptor (red) fluorophores for the smFRET_L1_ (above) and smFRET_L1-tRNA_ (below) signals depicted in GS1 of RC^Phe^. **(c)** Representative E_FRET_ versus time trajectories for smFRET_L1_ (left) and smFRET_L1-tRNA_ (right) at 298 K (above) and 310 K (below) for RC^Phe^. **(d)** Histograms of E_FRET_ values collected using smFRET_L1_ (left) and smFRET_L1-tRNA_ (right) for RC^Phe^ at five temperature points between 298 K and 310 K.

A very powerful framework for understanding the contributions of ligand interactions to the conformational dynamics of biomolecules is provided by free energy landscape theory (*31,32*). Using this framework, we define two ensembles of RC conformations corresponding to the initial and final states of the conformational change described above. We refer to these as Global States 1 and 2 (GS1 and GS2) and represent them as occupying two distinct local minima (*i*.*e*., wells) on a free energy landscape where each point on the landscape corresponds to the free energy of a unique RC conformation. Given the compound nature of conformational transitions between GS1 and GS2 (GS1→ GS2 and GS2→ GS1 transitions), these transitions encompass several large-scale structural rearrangements of the RC, including changes to: (*i*) the relative orientation between the small, 30S, and large, 50S, ribosomal subunits that make up the complete 70S ribosome; (*ii*) the position of the tRNA between two binding sites on the ribosome; and (*iii*) the location of a ribosomal structural element called the L1 stalk between two positions on the surface of the ribosome (*22*). On the free energy landscape, GS1→ GS2 and GS2→ GS1 transitions are represented as excursions of the RC over a higher free energy region of the landscape called the ‘transition state ensemble’ (TSE) that acts as a free energy barrier separating GS1 and GS2 (Fig. 1a). The height of this barrier (ΔG^‡^), which corresponds to the difference in free energy between GS1 or GS2 and the TSE, governs the rate of transition between GS1 and GS2. In the case of GS1→ GS2 and GS2→ GS1 transitions, smFRET (*10,18,25,26*) and cryogenic electron microscopy (cryoEM) (*33–35*) experiments have been used to measure the rate constants that directly yield ΔG^‡^ after analysis with an appropriate theoretical model, such as transition state theory (TST) (36). Of much more interest, but as yet unmeasured for these transitions or, to the best of our knowledge, for any comparable conformational transition in a biomolecular machine, are the energetic components of the ΔG^‡^, known as the activation enthalpy (ΔH^‡^) and activation entropy (ΔS^‡^). These parameters directly report on the physical and structural properties of the biomolecular machine that give rise to the barrier in the first place and, consequently, uniquely provide molecular insights into the mechanisms that ligands use to modulate the conformational dynamics of the machine (Fig. S2)

By analyzing how the rates of GS1→ GS2 and GS2→ GS1 transitions depend on temperature, we have resolved the free energy barriers of the RCs described above into their component ΔH^‡^s and ΔS^‡^s. Our measurements show that, while binding of tRNAs to the RC contributes to the barrier by remodeling intermolecular interactions within the RC (*i*.*e*., enthalpically) as well as by altering the structural flexibility of the RC and/or disorder of the surrounding solvent shell (*i*.*e*., entropically), it is the entropic component that modulates the dynamics of the RC in a manner that is conducive to overall ribosomal function. This tRNA-dependent entropic control of ribosome dynamics is the strategy that the ribosome has evolved to enable rapid GS1→ GS2 and GS2→ GS1 transitions while maintaining a tight interaction with the tRNA in both states. Since these considerations apply not just to ribosomes but to all biomolecular machines, we hypothesize that such ligand-dependent entropic control is a generalized mechanism for regulating the functional dynamics of such machines.

## RESULTS

### Experimental design

In this study, we used two previously characterized smFRET signals, each of which reports on a different aspect of the ribosomal structural changes comprising GS1→ GS2 and GS2→ GS1 transitions (*10,18*). The first follows the position of the L1 stalk from the ribosomal frame of reference (*18*) (smFRET_L1_), while the second reports on the relative distance between the L1 stalk and the ribosome-bound tRNA (*10*) (smFRET_L1-tRNA_) (Fig. 1b). The distance-dependent FRET efficiencies (E_FRET_s, defined as the fluorescence intensity of the FRET acceptor fluorophore normalized by the sum of the fluorescence intensities of the FRET donor and acceptor fluorophores) for these signals allowed us to follow the motions of the L1 stalk alone, and the combined motions of the L1 stalk and the tRNA, respectively, as the RCs transitioned between GS1 and GS2 (Fig 1c). Employing a novel, high precision, temperature-controlled, microfluidic flow-cell that we have previously developed for use in a single-molecule total internal reflection fluorescence (TIRF) microscope (*37*), we performed smFRET experiments using these signals at temperatures between 298 K and 310 K on a range of different RCs. These include RCs carrying a deacylated tRNA specific to phenylalanine (tRNA^Phe^) (RC^Phe^); carrying a deacylated tRNA specific to formylmethionine (tRNA^fMet^) (RC^fMet^); and lacking a tRNA altogether (RC^vacant^). tRNA^Phe^ belongs to the ‘elongator’ class of tRNAs (*25*), that are responsible for decoding the triplet-nucleotide codon sequence of the mRNA and consecutively adding the corresponding amino acids (in this case, phenylalanine) to the growing polypeptide chain during the elongation phase of translation. On the other hand, tRNA^fMet^ belongs to the ‘initiator’ class of tRNAs (*25*), that are responsible for decoding the mRNA start codon and directing the RC to assemble at the correct location on the mRNA during the initiation phase of translation. Thus, comparisons between the free energy landscapes of these different RCs allow us to investigate the unique energetic contributions made by tRNAs to facilitate different stages of translation (Fig. S1). To obtain the necessary information for those comparisons, we used TST to model the GS1→ GS2 and GS2→ GS1 transitions for each RC with temperature-dependent ΔG^‡^s (Fig. 1c and Methods) and subsequently calculated ΔH^‡^s and ΔS^‡^s from the temperature-independent and dependent components, respectively, of the ΔG^‡^s for each RC (*36*).

At temperatures near 310 K and above, the GS1→ GS2 and GS2→ GS1 transitions became too fast to be accurately detected by the electron-multiplying charge-coupled device (EMCCD) camera on our TIRF microscope (Fig. 1c), leading to potential errors in our estimates for rate constants (*37,38*). We therefore used a novel, time-resolution-enhancing, machine-learning algorithm that we call <u>*B*</u>ayesian <u>*I*</u>nference for the <u>*A*</u>nalysis of <u>*S*</u>ub-temporal-resolution <u>*D*</u>ata (BIASD) (*39*) to analyze the E_FRET_ *versus* time trajectories recorded for each RC. Briefly, BIASD analyzes the distribution of E_FRET_ values collected from an entire ensemble of RC molecules, and infers the rate constants for the forward and reverse transitions (in this case, the GS1→ GS2 and GS2→ GS1 transitions, respectively) that yield the observed E_FRET_ distribution. Because even transitions that are too fast to be detected by the EMCCD will lead to broadening of the E_FRET_ distribution (Fig. 1d, and Fig. S3), BIASD enables us to accurately estimate rate constants for even the fastest dynamics that we observe in our smFRET experiments. Applying BIASD across the entire temperature range for each dataset directly yielded the underlying ΔH^‡^s and ΔS^‡^s that are responsible for the observed GS1→ GS2 and GS2→ GS1 dynamics for each RC and each smFRET signal (see Methods).

### The mechanism of coupling distinct motions within RCs

One of the longstanding questions in the study of ribosome dynamics is whether, and to what extent, the motions of distal structural elements within the RC are coordinated. This is particularly the case for the motions of the two ribosomal subunits, the tRNA, and the L1 stalk, which altogether make up the compound conformational rearrangement at the heart of GS1→ GS2 and GS2→ GS1 transitions. Based on previous studies, we and others have hypothesized that these motions are allosterically coupled (*18,23,26,28,29*). and that the relative rotation of the ribosomal subunits, which requires remodeling of a large number of inter-subunit interactions, must be the slowest and, therefore, rate-governing step for all of the other motions (*26*). Contrasting with this, other studies have been interpreted as providing evidence against such coupling (*27*). In our measurements here, we found that the energetics of the GS1⇌GS2 equilibrium for both RC^Phe^ and RC^fMet^ are independent of the smFRET signal used to measure them. Specifically, we found that the ΔH^‡^ and ΔS^‡^ for both the GS1→ GS2 and GS2→ GS1 transitions measured using either smFRET_L1_ or smFRET_L1-tRNA_, which follow different aspects of the structural rearrangements between GS1 and GS2, are within experimental uncertainty for both RCs (Fig. 2a and Table S1). This observation provides the most direct and strongest evidence to date that the motions of the L1 stalk and the ribosome-bound tRNA within an RC are directly coupled and that the barriers which control the motions of these RC components must have the same underlying physical and structural basis, and thus arise from the same rate-governing step. The large ΔH^‡^s that we observe for the GS1→ GS2 and GS2→ GS1 barriers in all three RCs, including RC^vacant^, which lacks a bound tRNA (Fig. 3a and Table S1), suggest that this rate-governing step involves remodeling of a large number of intermolecular interactions. This is consistent with the large number of inter-subunit interactions that must be remodeled during the relative rotation of the subunits, providing strong evidence that whatever process it is that governs the rate of L1 stalk and tRNA motions during GS1→ GS2 and GS2→ GS1 transitions, it involves intersubunit rotation.

**Figure 2.**
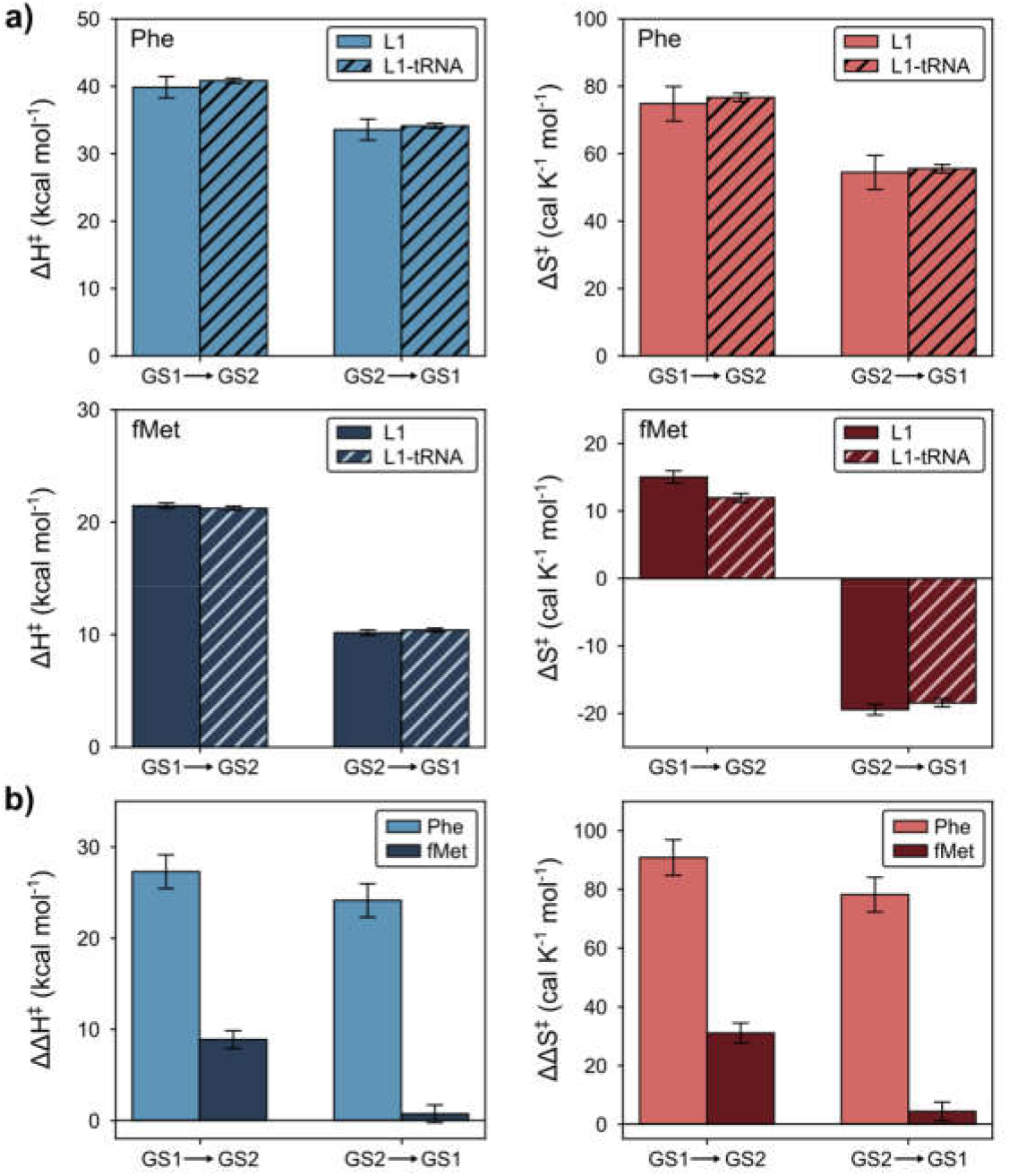
Activation parameters characterizing the GS1⇌ GS2 equilibrium in different RCs using multiple smFRET signals. **(a)** Comparisons of the activation enthalpies (ΔH^‡^, blue for RC^Phe^ and dark blue for RC^fMet^) and entropies (ΔS^‡^, red for RC^Phe^ and dark red for RC^fMet^) for smFRET_L1_ (solid) and smFRET_L1-tRNA_ (hatched). Error bars represent the standard deviations of the marginalized posterior distributions for the estimates (see Methods, and Table S1 for the number of individual molecules per dataset). **(b)** Relative activation enthalpies (ΔΔH^‡^, blue for RC^Phe^ and dark blue for RC^fMet^) and entropies (ΔΔS^‡^, red for RC^Phe^ and dark red for RC^fMet^) using RC^vacant^ as the common reference for smFRET_L1_. Error bars represent the standard deviations of the marginalized posterior distributions for the estimates (see Methods, and Table Sf1 for the number of individual molecules per dataset).

**Figure 3.**
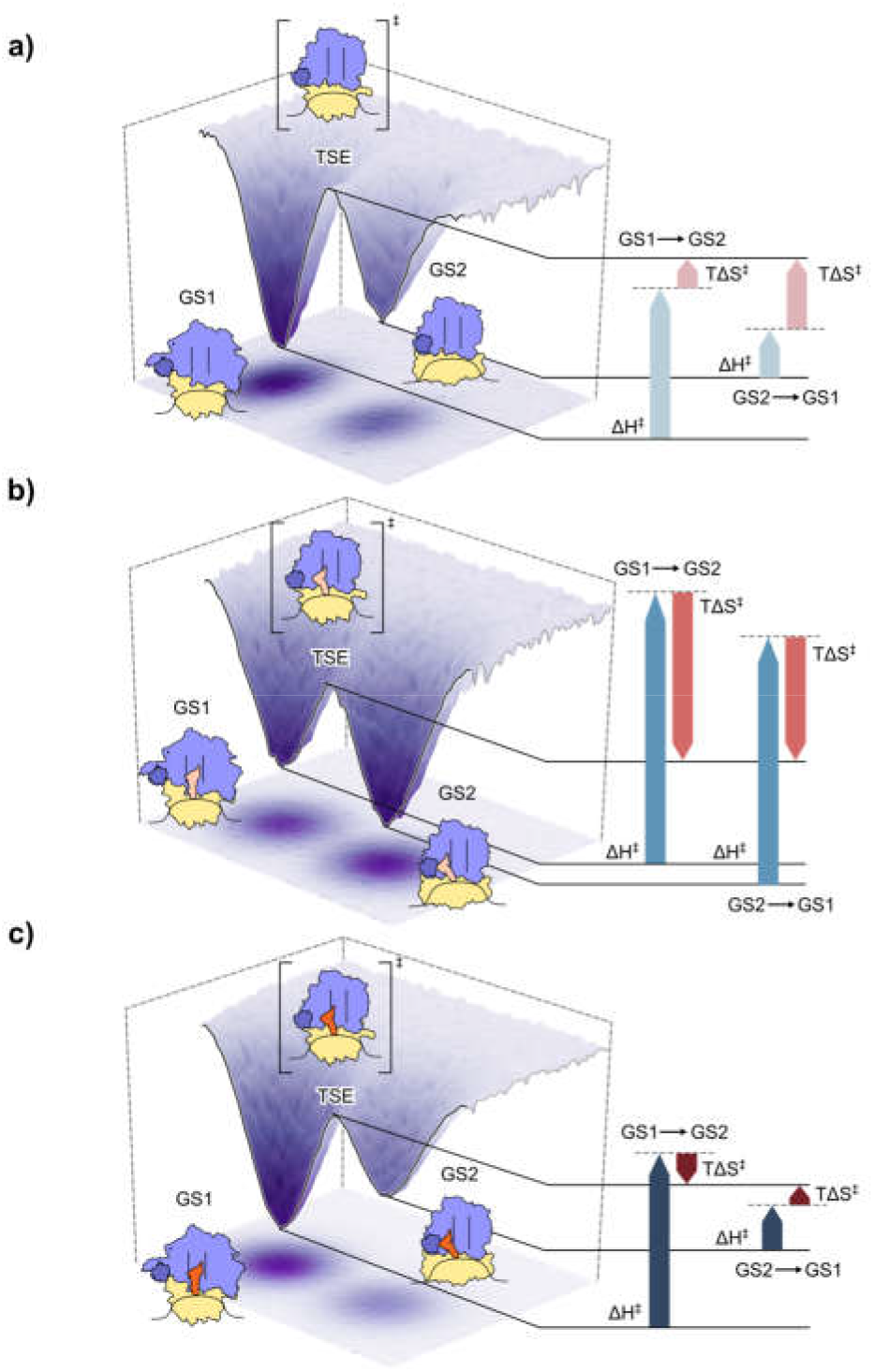
Enthalpic and entropic contributions to the free energy barriers separating GS1 and GS2 in different RCs. Structural cartoon representations of GS1, GS2, and the hypothetical TSE over the corresponding energy landscapes for **(a)** RC^vacant^, **(b)** RC^Phe^, and **(c)** RC^fMet^. The enthalpic (blue) and entropic (red) contributions to the free energy barriers separating GS1 and GS2 at a fixed temperature around 310K are shown for each RC. For each RC, the solid lines represent the free energies of GS1, GS2 and the TSE, while the dotted lines represent the free energies of the TSE if the corresponding free energy barriers had been solely enthalpic in nature. The TSE of RC^vacant^ illustrates disruption of the intersubunit interactions, while those for RC^Phe^ and RC^fMet^ shows the additional disruption of tRNA interactions to the binding sites on the large subunit that needs to occur along with the disruption of the intersubunit interactions.

Nonetheless, we also observe that the presence and identity of the ribosome-bound tRNA modulate the ΔH^‡^s and ΔS^‡^s of these barriers (Fig. 2a and Table S1). This leads us to the conclusion that the rate-governing step of the GS1→ GS2 and GS2→ GS1 transitions cannot be solely explained by the remodeling of inter-subunit interactions, and must include remodeling of tRNA-ribosome interactions. Specifically, the TSE for these transitions must involve the disruption of the interactions between the tRNA and its binding site in GS1 and/or GS2 that precede or be concurrent with the disruption of the inter-subunit interactions during intersubunit rotation. By necessitating this order of events, the architecture of the RC can couple the motion of the tRNAs with the internal rotational motion of the ribosome, which, in turn, also couples it to the motion of the L1 stalk (*26*), forming an allosteric network encompassing distal structural elements of the RC.

The findings we report here not only reveal that the motions of distal structural elements of the RC are coupled, but by uniquely providing information on the features of the TSE that control these dynamics (*i*.*e*., the ΔH^‡^s and ΔS^‡^s), they lead us to a mechanism for how coupling is achieved. The allosteric network formed by the motions of these structural elements also readily explains how changes made to any one element (*e*.*g*., the tRNA) can modulate the structural dynamics of the RC as a whole—a strategy which, most likely, was evolved to regulate the dynamics of the ribosome in a manner that aids it function (see below).

### tRNA-specific enthalpic penalties to RC dynamics

Beyond the coupling of motions within RCs, a further outstanding question in the field is how the specific interactions that particular tRNAs make to the ribosome modulate the dynamics of RCs. To address this question, we next compared the relative effects that different tRNAs have on the free energy barriers separating GS1 and GS2. Since the presence of ribosome-bound tRNA is the only difference between RC^vacant^ and the other RCs, the relative changes in ΔH^‡^ measured using smFRET_L1_ (*i*.*e*., ΔΔH^‡^_Phe_ = ΔH^‡^_Phe_ – ΔH^‡^_vacant_, and, similarly, ΔΔH^‡^_Phe_ = ΔH^‡^_Phe_ – ΔH^‡^_vacant_) should reveal the enthalpic effects a ribosome-bound tRNA has on the GS1→ GS2 and GS2→ GS1 transitions. We found that ΔΔH^‡^_Phe_ is positive for both the GS1→ GS2 and GS2→ GS1 transitions (27 kcal mol^−1^ and 25 kcal mol^−1^, respectively), demonstrating that significant tRNA^Phe^-specific interactions need to be remodeled for both of these transitions in RC^Phe^ (Fig. 2b and Fig. 3b). The ΔΔH^‡^_fMet_ values for both the GS1→ GS2 and GS2→ GS1 transitions, while still positive, are much lower (8 kcal mol^−1^ and 0.8 kcal mol^−1^, respectively), indicating that tRNA^fMet^-specific interactions with the ribosome are significantly weaker, at least at the sites that are remodeled during these transitions (Fig. 2b and Fig. 3c)

Strikingly, the very small ΔΔH^‡^_fMet_ value for the GS2→ GS1 transition (an order of magnitude less than the other ΔΔH^‡^ values) suggest that there are no significant tRNA^fMet^-ribosome interactions in GS2 that are remodeled upon the transition to GS1 or that these interactions are almost completely compensated by favorable remodeling of intramolecular tRNA^fMet^-tRNA^fMet^ interactions during the transition. Despite this, structural studies of RCs analogous to RC^Phe^ and RC^fMet^ show that, in GS2, both tRNA^Phe^ and tRNA^fMet^ exist in very similar conformations (*40,41*). Indeed, analysis of the positions of the tRNA and the L1 stalk (which forms part of the tRNA binding site in GS2) in these structures demonstrates that, in spite of slight positional differences, tRNA^Phe^ and tRNA^fMet^ both exist in close proximity to the L1 stalk in GS2 (Fig. S4). These structural observations agree with the very similar, high E_FRET_ values for GS2 that we observe in the smFRET_L1-tRNA_ signal for RC^Phe^ and RC^fMet^ (0.81 and 0.80, respectively). Collectively, the small ΔΔH^‡^_fMet_ value for the GS2→ GS1 transition and relative positioning of L1 stalk and tRNA^fMet^ in GS2 leads us to conclude that the motion of tRNAs within RCs occurs as a consequence of the architecture of the ribosome, regardless of any stabilizing interactions that may or may not be formed between ribosomal elements and the tRNA in either GS1 or GS2.

We should note at this point that, while the enthalpic differences between the RCs that we observe may in part originate from the remodeling of solvent interactions at the tRNA-binding sites of the RCs, the significant dependence of these ΔΔH^‡^s on the identity of the bound tRNA strongly suggest that the major contributions to the enthalpic differences derive from the remodeling of intermolecular tRNA-RC and intramolecular tRNA-tRNA interactions. Regardless of the molecular origin of these effects, we hypothesize that RCs have evolved to harness such enthalpic differences as a way to allosterically modulate their conformational dynamics through perturbations from only a single type of molecular component—in this case, the tRNAs. Unique tRNAs might then have evolved to form different interactions at the tRNA-ribosome interface with the goal of allosterically modulating the functional dynamics of the entire ribosome.

### tRNA-induced entropic compensation of enthalpic penalties to RC dynamics

All things kept equal, the enthalpic penalties described above should have increased the activation barriers, and thus decreased the rate constants of the GS1→ GS2 and GS2→ GS1 transitions in tRNA-bound RCs relative to RC^vacant^. Instead, however, we found that these rate constants were similar to or larger in the tRNA-bound RCs than in RC^vacant^ (Fig. S5). This was driven by the fact the ΔΔS^‡^ (defined similar to ΔΔH^‡^ above) for the GS1→ GS2 and GS2→ GS1 transitions were also positive. At the temperatures used in our measurements (and especially around 310 K, which is the optimal growth temperature for *E. coli* (*42*)), the ΔΔS^‡^_Phe_ s for the GS1→ GS2 and GS2→ GS1 transitions (91 cal K^−1^ mol^−1^ and 77 cal K^−1^ mol^−1^, respectively) more than compensate for the enthalpic penalties observed for RC^Phe^ in comparison to RC^vacant^, leading to faster dynamics at the higher temperatures (around 310 K) (Fig. 2b and Fig. 3b). The same is true for the ΔΔS^‡^_fMet_ values for the GS1→ GS2 and GS2→ GS1 transitions (31 cal K^−1^ mol^−1^ and 4 cal K^−1^ mol^−1^, respectively), even if they are smaller in comparison to RC^Phe^ (Fig. 2b and Fig. 3c). In fact, in the absence of any significant opposing enthalpic penalty, the small ΔΔS^‡^_fMet_ for the GS2→ GS1 transitions is enough to increase the corresponding rate constant by nearly three-fold over that of RC^vacant^ over the entire range of temperature, making it the fastest transition we observe in our study (Fig. S5).

Surprisingly, we find that the ΔΔS^‡^ values we measure are largely correlated with their corresponding ΔΔH^‡^ values. While the reason for this enthalpy-entropy compensation is not immediately obvious, we hypothesize this is necessary to not overcompensate the tRNA-induced enthalpic penalties for these transitions. The GS1→ GS2 and GS2→ GS1 transitions are part of a large number of structural rearrangements that RCs must undergo during translation. All of these rearrangements need to occur within a very specific kinetic regime for translation to occur as rapidly as possible while maintaining the integrity of the process. Specifically, while slowing down GS1→ GS2 and GS2→ GS1 transitions as a result of uncompensated tRNA-induced enthalpic penalties could hinder translation and therefore be detrimental to cellular fitness, speeding up these transitions too much *via* compensatory entropic modulations might result in inaccurate tRNA or ribosome movements that could be deleterious to translation (*e*.*g*., slipping of the RC on the mRNA). The need to optimize this trade-off between speed and accuracy would explain the enthalpy-entropy compensation that we observe and highlight the evolutionary pressures that underly these energetic modulations of ribosome dynamics.

Similar to the case for the ΔΔH^‡^s above, the ΔΔS^‡^s that we observe may arise either from modulating the available conformational entropy of the RCs themselves or from a change in the ordering of water molecules, metal cations, and/or polyamines around the RCs. Indeed, the contributions from these sources of entropy are not mutually exclusive and further studies are required to parse out the effects of conformational and solvent entropies to these free energy barriers. Regardless of the relative contributions of either source, we note that it is these ligand-dependent entropic modulations, and not the corresponding enthalpic penalties, which drive the rates of GS1→ GS2 and GS2→ GS1 transitions in a manner that allows protein synthesis to take place rapidly, but accurately.

### The net effect of entropic modulations to the dynamics of different tRNA-bound RCs

Our measurements show that both tRNA^Phe^ and tRNA^fMet^ employ the strategy of entropic modulation to overcome specific tRNA-induced enthalpic penalties and drive the functional dynamics of their corresponding RCs. However, we see that the net effects of these modulations are markedly different for the respective RCs (Fig. 3). In RC^Phe^, the effect of the tRNA^Phe^-specific entropic modulations leads to a larger increase in the GS1→ GS2 transition rate than the GS2→ GS1 transition rate, biasing the GS1⇌GS2 equilibrium towards GS2. In contrast, tRNA^fMet^-specific entropic modulations in RC^fMet^ lead to a more significant increase in the GS2→ GS1 transition rate over the GS1→ GS2 transition rate, which biases the GS1⇌GS2 equilibrium towards GS1. In the context of the specific steps of translation these tRNAs are involved in (*i*.*e*., elongation *versus* initiation, respectively) (Fig. S1), it is clear that these entropic modulations bias the GS1⇌GS2 equilibrium towards the respective state responsible for productive, forward progression through the translation cycle (*i*.*e*., towards GS2 for elongation and towards GS1 for initiation).

Interestingly, comparing the GS2 structures of an RC analogous to RC^Phe^ (*40*) and an RC analogous to RC^Phe^, but carrying a different elongator tRNA, tRNA^Met^ (the elongator tRNA specific to methionine), (RC^Met^) (*43*), shows that the L1 stalk-tRNA interface in both structures are more similar to each other than to the one present in RC^fMet^ (Figs. S4 and S6). Given the similarities between the structures of RCs carrying different elongator tRNAs, and, indeed, the similarities in the rates of GS1→ GS2 and GS2→ GS1 transitions in such RCs (*25*), we hypothesize that the tRNA^Phe^-induced entropic modulation of RC dynamics we observe generalizes to other elongator tRNAs and collectively serve to speed up elongation. Analogously, we hypothesize that the tRNA^fMet^-induced entropic modulation of RC dynamics we observe is used to speed up initiation instead. Taken together, our findings strongly suggest that tRNAs and ribosomes have co-evolved to utilize a complex interplay of ligand-induced enthalpic and entropic modulation to control the conformational dynamics of the entire biomolecular complex in ways that facilitate the overall process of translation.

We note that, while our experiments focus on the entropic and enthalpic effects of tRNA binding to RC dynamics, a multitude of protein translation factors, RNA accessory factors, and mRNA structural elements interact with the RC during translation. The binding of many of these factors has also been shown to modulate RC dynamics in a manner that facilitates the specific steps of translation that each is involved in (*18,23,24*). While the entropic and enthalpic contributions to the free energy barriers underlying the dynamics of such factor-bound RCs have not yet been characterized, our results readily point to a mechanism for the compensation of the enthalpic penalties of factor binding to these RC using similar entropic modulation strategies.

## DISCUSSION

Our experimental results reveal the existence of tRNA-induced entropic modulation of RC dynamics and uncovers their role in kinetically facilitating those dynamics. The unique data provided by our temperature-dependent, single-molecule experiments have allowed us to characterize the thermodynamics of the TSE governing RC dynamics in a manner that is inaccessible to structural studies or kinetics studies performed at a single temperature (*36, 44*). For example, the increased ΔS^‡^s for the GS1→ GS2 and GS2→ GS1 transitions in RC^Phe^ and RC^fMet^ suggest that the TSEs in these RCs are relatively more flexible or disordered relative to that in RC^vacant^ (Fig. S2c). We hypothesize that it is this increase in the structural flexibility of the TSEs that leads to the faster rates of GS1→ GS2 and GS2→ GS1 transitions in the tRNA-bound RCs. In terms of free energies, the increase in structural flexibility corresponds to an increase in the number of microstates that have sufficiently low free energy to enable transitions between GS1 and GS2, thereby leading to an expansion of the TSE. This expansion provides a greater number of possible paths across the free energy barrier separating GS1 and GS2, consequently increasing the rates of transitions between the two states (Fig. 4). We contrast this entropically driven increase in transition rates with an enthalpically driven mechanism commonly encountered in small enzyme kinetics (*36*), where the formation of stabilizing interactions in the TSE lowers the free energy barrier between two states. We note that there is no such net change in favorable interactions in the TSEs of the RCs that we studied. The stabilizing interactions we do encounter are net unfavorable for the transitions between GS1 and GS2, and therefore need to be compensated for using entropic modulations. These differences in the relative roles of enthalpic and entropic contributions between the dynamics commonly seen in small proteins and the dynamics we see in the ribosome suggest that our understanding of the physical principles controlling small enzyme dynamics need not be readily translated to the dynamics observed in these machines. Our results, thus, highlight the need for further theoretical and experimental investigations of the dynamics of the ribosome and similar biomolecular machines.

**Figure 4.**
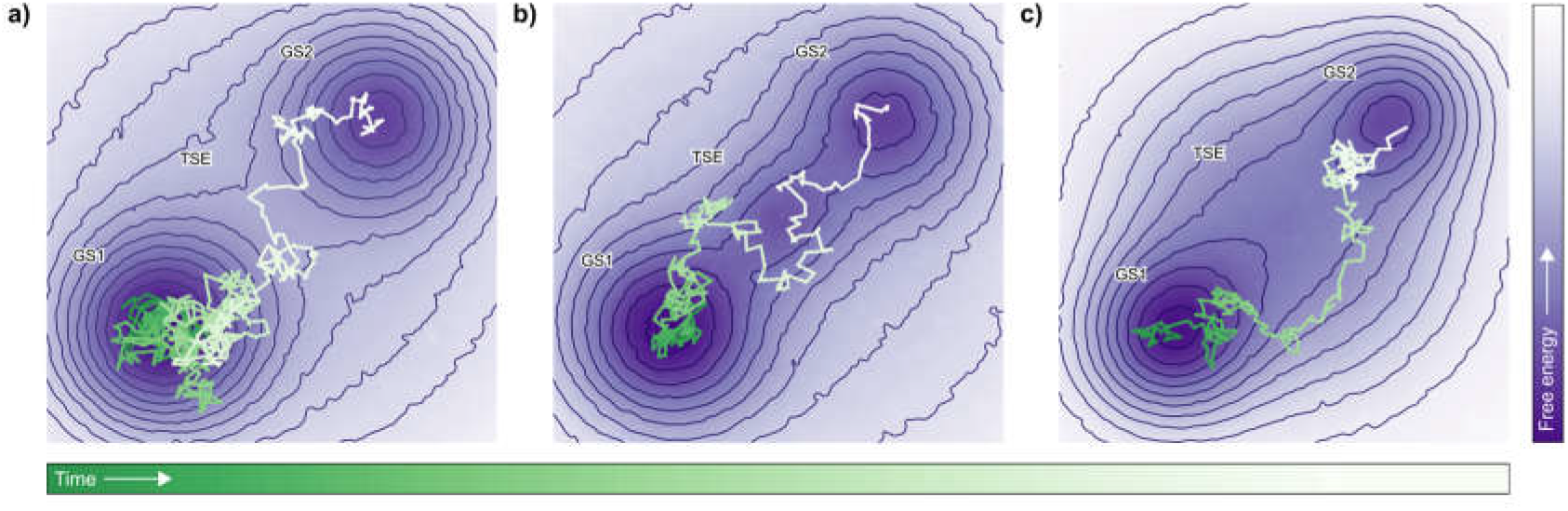
Entropic modulation increases the rate of structural transitions. **(a)** A simulated random walk trajectory (green to white as a function of increasing time) consisting of 725 steps is shown on a contour plot of a hypothetical free energy landscape (dark purple to light purple as a function of increasing free energy) consisting of the two conformational states, GS1 and GS2 (dark purple wells) separated by a TSE on the free energy barrier (light purple region of high free energy separating the two wells). The random walk moves from GS1 to GS2 via the TSE. **(b)** Enthalpic stabilization of the TSE causes a lowering of the barrier between GS1 and GS2 that, at an extreme, can result in the creation of an intermediate state. Lowering of the barrier in this way allows a shorter simulated walk of 290 steps to traverse the landscape from GS1 to GS2, demonstrating the resultant increase in the rate of transitions. **(c)** Entropic modulation of the free energy landscapes, represented by increasing the size of the TSE, widens the barrier, allowing more possible trajectories to successfully cross the barrier. The simulated trajectory of 253 steps shown here demonstrates how expanding the TSE leads to more possible paths across the barrier and an associated increase in the rate of transitions. Such a trajectory would not have been possible in the narrower barrier in (a).

tRNA-induced entropic modulation of RC dynamics appears to be a mechanistic strategy used to overcome the enthalpic penalties associated with tRNA binding to the ribosome while simultaneously optimizing the dynamics of the RC such that it can successfully navigate the speed/accuracy trade-off that is inherent to mechanical processes such as translation. Additionally, by evolving an architecture that enables the allosteric coupling of multiple distal structural elements, RCs can utilize perturbations in a single component (in this case, the tRNA) to modulate the dynamics of the entire biomolecular complex. Moreover, by utilizing perturbations from ligands like tRNAs, which are modular and can be readily replaced in different contexts, RCs can differentially bias the same set of dynamics for multiple purposes at different phases of translation. Indeed, the generality of the constraints that these ligand-induced entropic modulations have evolved to overcome leads us to hypothesize that similar modulations must also play a major role in the regulation of the functional dynamics of other multi-component biomolecular machines. Given their size and complexity, the dynamics of all such machines must be associated with significant remodeling of inter- and intramolecular interactions, especially in the presence of ligands which need to be tightly bound and moved across distal sites in the complex. The work we present here shows that when constraints exist on the possible number of ways in which the enthalpy of a biomolecular machine can be modulated, then biomolecular machines can evolve coherent strategies to instead utilize entropy to drive and regulate their conformational dynamics, and thus, their functions.

## CONCLUSIONS

The use of enthalpy-entropy compensation has been previously observed in a range of biomolecular systems, including the folding of proteins (*45*), the recognition of ligands (*46*), and the kinetics of enzyme reactions (*47*). In this work, we show that the function and regulation of biomolecular machines also rely on enthalpy-entropy compensation and provide a detailed characterization of this compensation in the operation of a paradigmatic biomolecular machine. Given the ubiquity and diversity of biological processes performed by such large biomolecular complexes, we predict that the mechanisms for utilization of ligand-dependent entropic control of free energy landscapes that we describe here will be a key concept that furthers our understanding of the workings of natural biomolecular machines and form an important design principle for the development of synthetic, bio-mimetic molecular machines.

## MATERIALS AND METHODS

### Preparation of purified fluorophore-labeled and unlabeled translational components

Fluorophore-labeled ribosomes and tRNAs were prepared following published protocols (*12*). Briefly, a single-cysteine (Cys), *E. coli* ribosomal protein uL1 variant, uL1_T202C_, and a single-Cys, *E. coli* ribosomal protein bL9 variant, bL9_Q18C_, were designed, overexpressed, purified, and labeled with maleimide-activated Cy5 or Cy3, respectively. Cy5-labeled uL1_T202C_ and Cy3-labeled bL9_Q18C_ were then reconstituted into large ribosomal subunits lacking both wild-type uL1 and bL9 that had been purified from a uL1-bL9, double-deletion, *E. coli* strain, using multiple sucrose density gradient ultracentrifugation steps, to generate Cy5-Cy3, dual-labeled large subunits. Similarly, Cy5-labeled uL1_T202C_ was reconstituted into large ribosomal subunits lacking wild-type uL1 that had been purified from a uL1-deletion *E. coli* strain to generate Cy5-labeled large subunits. *E. coli* tRNA^fMet^ and tRNA^Phe^ were labeled with Cy3-maleimide or N-hydroxysuccinimidyl (NHS) ester-activated Cy3 at the naturally post-transcriptionally modified 4-thiouridine at nucleotide position 8 (s^4^U8) and 3-(3-amino-3-carboxypropyl)uridine at nucleotide position 47 (acp^3^U47), respectively, to generate (Cy3)tRNA^fMet^ and (Cy3)tRNA^Phe^.

Unlabeled components were purified following published protocols (*12*). Briefly, unlabeled small ribosomal subunits were purified using multiple sucrose density gradient ultracentrifugation steps. Unlabeled translation factors, specifically initiation factors 1, 2 and 3 (IF1, IF2, IF3), and elongation factors Tu and G (EF-Tu and EF-G), were purified using affinity chromatography, followed by subsequent size exclusion chromatography and/or cation exchange chromatography, as previously described (*12*). tRNA^fMet^, (Cy3)tRNA^fMet^, tRNA^Phe^, and (Cy3)tRNA^Phe^ were aminoacylated with the corresponding amino acids and aminoacyl tRNA synthetases, and Met-tRNA^fMet^ and Met-(Cy3)tRNA^fMet^ were formylated using formylmethionyl-tRNA formyltransferase, as previously described (*12*).

### Preparation of RC^vacant^, RC^Phe^, and RC^fMet^

RCs containing tRNAs were enzymatically assembled *in vitro* in the manner which had been previously described for temperature dependent smFRET measurements of RC^Phe^ (*37*). Briefly, RC^fMet^ were prepared by enzymatically initiating unlabeled small subunits and large subunits (Cy5-Cy3 dual-labeled for smFRET_L1-tRNA_ or Cy5 labeled for smFRET_L1_) at a start codon on a 5’-biotinylated mRNA with fMet-tRNA^fMet^ (for smFRET_L1_) or fMet-(Cy3)tRNA^fMet^ (for smFRET_L1-tRNA_) and translation initiation factors 1, 2, and 3. RC^Phe^ was prepared by enzymatically elongating RC^fMet^ prepared as described above using unlabeled fMet-tRNA^fMet^ and Cy5-Cy3 dual-labeled large subunits (for smFRET_L1_) or Cy5 labeled large subunits (for smFRET_L1-tRNA_), with Phe-tRNA^Phe^ or Phe-(Cy3)tRNA^Phe^ respectively, delivered by translation elongation factor Tu and subsequently translocated by translation elongation factor G. RC^vacant^ were assembled non-enzymatically according to a previously described protocol (*26*). Briefly, this reaction was conducted in two steps: (i) incubating unlabeled small subunits with 5’-biotinylated mRNA in RC assembly buffer [50 mM Tris-Cl (pH_25°C_ = 7.5), 70 mM NH_4_Cl, 30 mM KCl, and 6 mM βME] containing 20 mM MgCl_2_, for 10 minutes at 37 °C, (ii) adding Cy5-Cy3 dual-labeled large subunits to the above reaction and incubating the resulting sample at 37 °C for 20 minutes. All RCs were purified using sucrose density gradient ultracentrifugation.

### Fabrication, assembly, calibration, and performance of temperature-controlled microfluidic flowcells

The fabrication, assembly, calibration, and performance of the temperature-controlled microfluidic flowcells used here for smFRET imaging has been previously described (*37,48*). Briefly, each flowcell consists of a set of five parallel microchannels that are sandwiched between a quartz microscope slide (G. Finkenbeiner) and a borosilicate glass coverslip (VWR). Prior to assembling the flowcell, the slide was passivated against non-specific binding of biomolecules by derivatizing the cleaned and aminosilanized (Vectabond, Vector Labs) quartz surface with a mixture of NHS ester-activated polyethylene glycol (NHS-PEG) and dilute NHS-PEG-biotin. Likewise, prior to assembling the flowcell, the coverslip was microfabricated in order to integrate thermal control elements. These elements included thin-film resistive microheaters that were distributed evenly across the length of each microchannel for uniform heating as well as thin-film resistive temperature sensors that were located along the long side and off-center of each microchannel to accurately probe the temperature in real time. Flowcells were then assembled by placing ∼1 mm wide strips of double-sided tape on each side of each microchannel on the slide and affixing the coverslip on top of the double-sided tape-containing face of the slide such that the fabricated face of the coverslip is positioned inside of the resulting flowcell. Subsequently, the sides of the flowcell were sealed with epoxy. The temperature sensor in each microchannel was then calibrated by characterizing the linear relationship between resistance and temperature such that the microchannel temperature could be accurately and precisely determined by measuring the temperature sensor resistance. Further performance characterization of the resulting temperature-controlled microchannels demonstrated that closed-loop control of the on-chip heaters and temperature sensors could accurately maintain the setpoint temperature to a precision of ± 0.01°C.

### smFRET experimental conditions

As previously described in Wang, *et al*. (*37*), experiments were carried out in Tris Polymix Buffer [50 mM tris(hydroxymethyl)aminomethane acetate (Tris-OAc) (pH_25°C_ = 7.0), 100 mM potassium chloride (KCl), 5 mM ammonium acetate (NH_4_OAc), 0.5 mM calcium acetate (Ca(OAc)_2_), 0.1 mM ethylenediaminetetraacetic acid (EDTA), 10 mM 2-mercaptoethanol (βME), 5 mM putrescine dihydrochloride, and 1 mM spermidine, free base] at 15mM magnesium acetate (Mg(OAc)_2_) (*12*), supplemented with an oxygen-scavenging system (300 μg/mL glucose oxidase (Sigma), 40 μg/mL catalase (Sigma), and 1% β-D-glucose), and a triplet-state quencher cocktail (1 mM 1,3,5,7-cyclooctatetraene (Aldrich), and 1 mM p-nitrobenzyl alcohol (Fluka)) (*12*). The biotin-functionalized RCs were flown into the microscope flowcell and surface immobilized utilizing biotin-streptavidin-biotin interactions. All RCs containing a peptide attached to the tRNA (RC^fMet^ and RC^Phe^) were incubated in 1 mM puromycin in Tris Polymix Buffer at room temperature for 5 minutes prior to imaging. This caused hydrolysis and release of the tRNA-bound nascent peptide, leading to the formation of RCs containing only a deacylated tRNA.

### smFRET imaging using total internal reflection fluorescence microscopy

As previously described in Wang, *et al*. (*37*) for the smFRET_L1-tRNA_ signal, imaging was performed with a laboratory-built, wide-field, prism-based total internal reflection fluorescence (TIRF) microscope. A 532 nm diode-pumped solid-state laser (CrystaLaser, Inc.) was used as an excitation source, and a 512 pixel × 512 pixel electron-multiplying charge-coupled-device camera (EMCCD) (Cascade II 512, Photometerics, Inc.) was used as a detector. For smFRET_L1-tRNA_ experiments, the excitation density was 32 W cm^−2^ (the excitation area is estimated to be 0.05 mm^2^), and the acquisition rate for the EMCCD was 20 frames sec^−1^. For smFRET_L1_ experiments, the experimental setup was nearly identical to the one described above and previously (*37*). The major experimental differences were an excitation density of 22 W cm^−1^, and an EMCCD acquisition rate of 10 frames sec^−1^. Experiments for all RCs were conducted at five temperatures, 25 °C, 28 °C, 31 °C, 34 °C, and 37 °C (or 298 K, 301 K, 304 K, 307 K, and 310 K). The temperature inside a designated microchannel for single-molecule measurements was maintained at the setpoint using closed-loop control of the on-chip heaters and temperature sensors to within an accuracy of ± 0.01 °C (*37*).

Pairs of Cy3 and Cy5 fluorophores corresponding to individual RCs were co-localized by aligning the fields-of-view containing the fluorescence emissions of hundreds of individual Cy3 and Cy5 fluorophores using an Iterative Closest Point (ICP) algorithm (*49*) to find the best affine transform between the two views. The fluorescence intensities of the co-localized Cy3 and Cy5 fluorophores were subsequently fit to 2D Gaussian point spread functions and extracted from the movies to generate raw Cy3 and Cy5 fluorescence intensity *versus* time trajectories (intensity trajectories), which were subsequently corrected using a 5.5% Cy3-to-Cy5 bleedthrough parameter. The corrected intensity trajectories were transformed into E_FRET_ trajectories by calculating E_FRET_ = I_Cy5_ /(I_Cy3_ + I_Cy5_) at each time point, where I_Cy3_ and I_Cy5_ correspond to the intensity for the Cy3 and Cy5 fluorophore channels, respectively. Outlying E_FRET_ datapoints arising from estimates of E_FRET_ made in the absence of sufficient Cy3 or Cy5 fluorescence (where E_FRET_ > 2.0 or where E_FRET_ < −1.0, respectively) were removed. Only E_FRET_ trajectories exhibiting the complete loss of Cy3 and/or Cy5 fluorescence in a single step, which likely occurs due photobleaching and/or photoblinking of the fluorophores and can be distinguished from decreased, but still non-zero levels of fluorescence caused by conformation-dependent changes in FRET, were selected for analysis. The E_FRET_ trajectories were truncated at the first timepoint prior to the timepoint at which either the Cy3 or Cy5 fluorophore photobleached. In E_FRET_ trajectories where photoblinking events occurred, only a single section of each E_FRET_ trajectory, corresponding to the time period before the first photoblinking event, between two photoblinking events, or between a photoblinking event and the photobleaching event, was selected for further analysis.

### Estimation of activation parameters using BIASD

The selected E_FRET_ *versus* time trajectories for each smFRET signal at every temperature point T (*T* = 298, 301, 304, 307, 310 K) for each RC were concatenated into a single one-dimensional E_FRET_ data vector ({*d(T*)}) consisting of the signals from each individual RC molecule of that specific type at the given temperature. The resulting set of vectors ({d}, consisting of individual {*d(T*)}’s) for each RC was analyzed using a global BIASD (*39*) algorithm. In this analysis, the {*d*} for each RC and each smFRET signal was modeled using seven temperature-independent parameters: two E_FRET_ means (*ϵ*_*GS1*_ or *ϵ*_*GS2*_) corresponding to the conformations GS1 and GS2 for each specific RC respectively, an E_FRET_ noise parameter (*σ*) describing the noise in the E_FRET_ signal in both conformational states, and four activation parameters (*ΔH*^‡^_(GS1→ GS2)_ and *ΔH*^*‡*^_(GS1→ GS2),_ and *ΔH*^*‡*^_(GS2→ GS1),_ and *ΔH*^*‡*^_(GS2→ GS1)_describing the free energy barriers for the GS1→ GS2 and GS2→ GS1 transitions respectively. The rates of interconversion between the two states, *k*_*GS1→ GS2*_ and *k*_*GS2→ GS1*_, that were observed in each {*d(T*)} were determined using the respective activation parameters according to TST (*36*) with the equation:

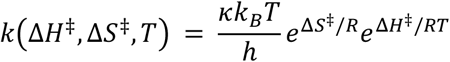

where *k*_*B*_ is the Boltzmann constant, *h* is the Planck constant, *R* is the gas constant, *T* is the absolute temperature at which the rate constant is observed, and *κ* is a correction factor which is set to 1 (*36*). This model was then used to explain the data {*d*} for each RC using Bayesian inference, Briefly, the log likelihood function used in this case was

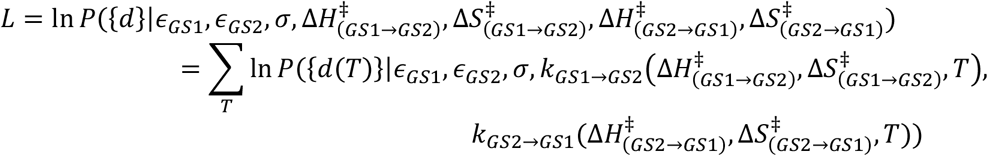

where ln *P*({*d*}|*ϵ*_*GS1*_, *ϵ*_*GS2*_, *σ, k*_*GS1→ GS2*_, *k*_*GS2→ GS1*_) is the BIASD log-likelihood function as has been derived previously (*39*) and {*T*} is the set of experimental temperatures from 298 K to 310 K. The prior distributions used can be found in the Table S3. The posterior probability distribution for each RC dataset, defined here as the product of the above likelihood and prior distributions (assuming the evidence is a constant, since only a single model is used) (*50*), was sampled using an affine-invariant Markov Chain Monte Carlo (MCMC) method, *emcee* (*51,52*). The sampling process was carried out in two steps. First, for each dataset, the MCMC method was initialized using 1000 walkers and 3000 steps. The final walker positions after 3000 steps were then sorted by posterior probability. The worst 500 positions were then discarded, and the remaining positions were used to initialize 500 new walkers, thereby eliminating sampling artefacts caused by improper initialization. In the second steps, these 500 walkers were employed to sample a further 6000 steps, the first 1000 of which were discarded to burn the chains. Uncorrelated samples were chosen from the remaining steps based on the maximum parameter correlation time (∼100 steps), and these uncorrelated samples were used to calculate the means and standard deviations of the marginalized posterior probability distributions for each RC dataset.

The analysis of the RC^Phe^ smFRET_L1_ data showed allowed us to specify mean E_FRET_ states for GS1 and GS2 with a much greater precision than could be previously achieved (*18,26*). The more precise posterior distributions for the mean E_FRET_ states were leveraged to specify tighter priors for the remaining two RCs for the smFRET_L1_ signal (*50*).

## ACKNOWLEDGEMENTS

We thank Dr. Paul Whitford (Northeastern University) for valuable comments on the manuscript. This work was supported by funds to R.L.G. from the National Institutes of Health (R01 GM084288, R01 GM137608, and R01 GM136960) and the National Science Foundation (MCB 0644262). C.D.K. was supported by funds from the U.S. Department of Energy Office of Science Graduate Fellowship Program (DE-AC05-06OR23100) and Columbia University’s NIH Training Program in Molecular Biophysics (T32-GM008281).

## DATA AND MATERIALS AVAILABILTY

The processed smFRET data, in the form of the 1-dimensional data vectors which may be readily used by our BIASD code, is stored in the compressed HDF5 file format in the experimental data directory of the Github repository (https://github.com/KorakRay/biasd-tst), along with additional information on the exact data structure used for storage. The unprocessed TIFF images of the smFRET experiments could not be stored in the same repository due to file size considerations, and are available from the corresponding author upon request. The code used in this study to analyze the raw TIRF movies and generate processed smFRET datasets is part of a custom Python package, vbSCOPE, that is available from R.L.G. upon request. The updated version of BIASD used in this work, which enables global analysis across multiple datasets, in this case at multiple temperatures, can be found in a Github repository (https://github.com/KorakRay/biasd-tst).

## Author Contributions

K.K.R., C.D.K., and R.L.G. designed the research; J.F. performed the smFRET experiments and collected the data; B.W. and Q.L. designed the temperature-controlled microfluidic flowcells; B.W. fabricated and calibrated the temperature-controlled microfluidic flowcells; K.K.R. analysed the data; K.K.R, C.D.K., and R.L.G. wrote the manuscript; all authors approved the final version of the manuscript.

## Competing Interest Statement

The authors declare no conflicts of interest.

## SUPPLEMENTARY MATERIALS

**Figure S1.**
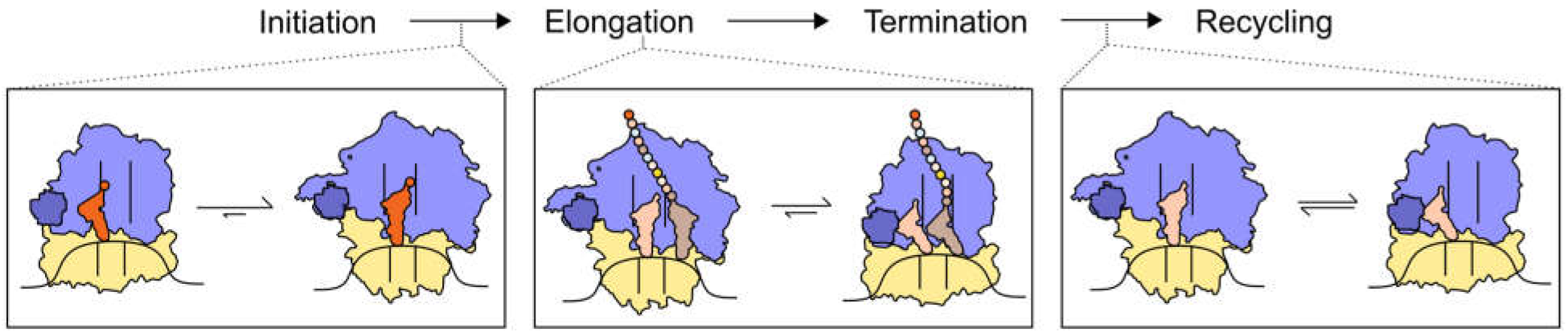
GS1⇌ GS2 or analogous equilibria in the translation pathway. Schematic of the four major stages of ribosome-catalyzed protein synthesis (top) and the role that regulation of the GS1⇌ GS2 equilibrium (or analogous equilibria between RC global states similar to GS1 and GS2) plays along this reaction pathway (bottom).

**Figure S2.**
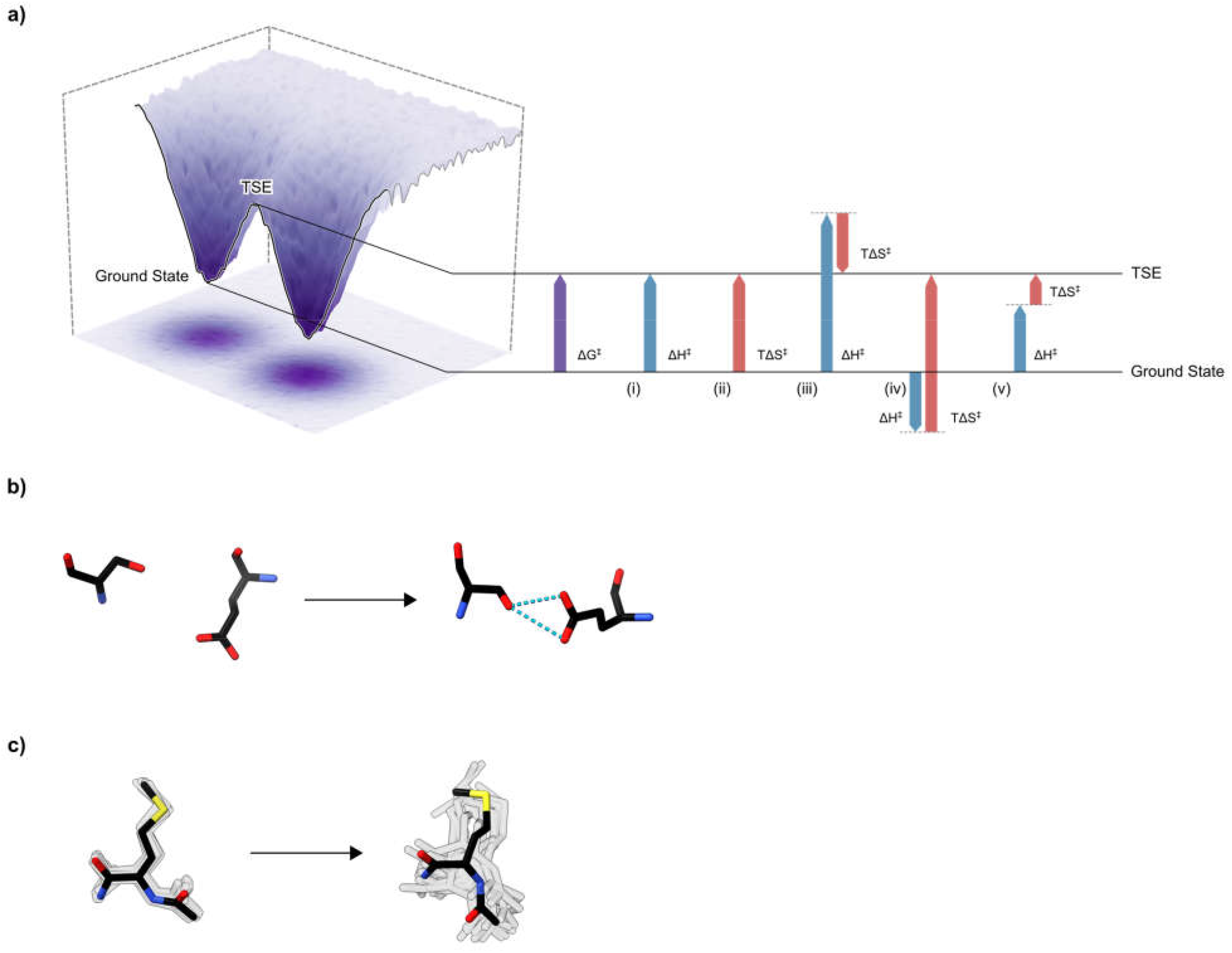
Free energy landscapes and their mechanistic implications. (**a**) A hypothetical free energy (*ΔG*^*‡*^) (purple) barrier and the combinations of enthalpic (*ΔH*^*‡*^) (blue) and entropic (*ΔS*^*‡*^) (red) components capable of comprising that barrier. Specifically, the barrier may be comprised of components that are (i) purely enthalpically unfavorable (ii) purely entropically unfavorable, (iii) enthalpically unfavorable-entropically favorable, (iv) enthalpically favorable-entropically unfavorable, or (v) enthalpically unfavorable-entropically unfavorable. In cases iii-v, the dotted line shows the height of the barrier if enthalpy was the only contributing component. Because the enthalpic component reports on remodeling of inter- and intramolecular interactions, and the entropic component reports on changes in the number of microstates (interpreted as modulating structural flexibility of the system) during the transition, each of i-v shows a different structural and physical mechanism underlying the same free energy barrier. **(b-c)** Representative structural interpretations of **(b)** enthalpically and **(c)** entropically favorable processes. The enthalpically favorable process leads to the formation of intermolecular interactions (in this case, hydrogen bonds) between the depicted serine and glutamate residues. The entropically favorable process leads to an increase in the number of microstates (in this case, conformational rotamers) of the depicted methionine residue, thereby making it more flexible. For completeness, we note that the reverse processes (i.e., breaking of the hydrogen bonds between the serine and glutamate residues and a decrease in the number of conformational rotamers of the methionine) would constitute **(b)** enthalpically and **(c)** entropically unfavorable processes.

**Figure S3.**
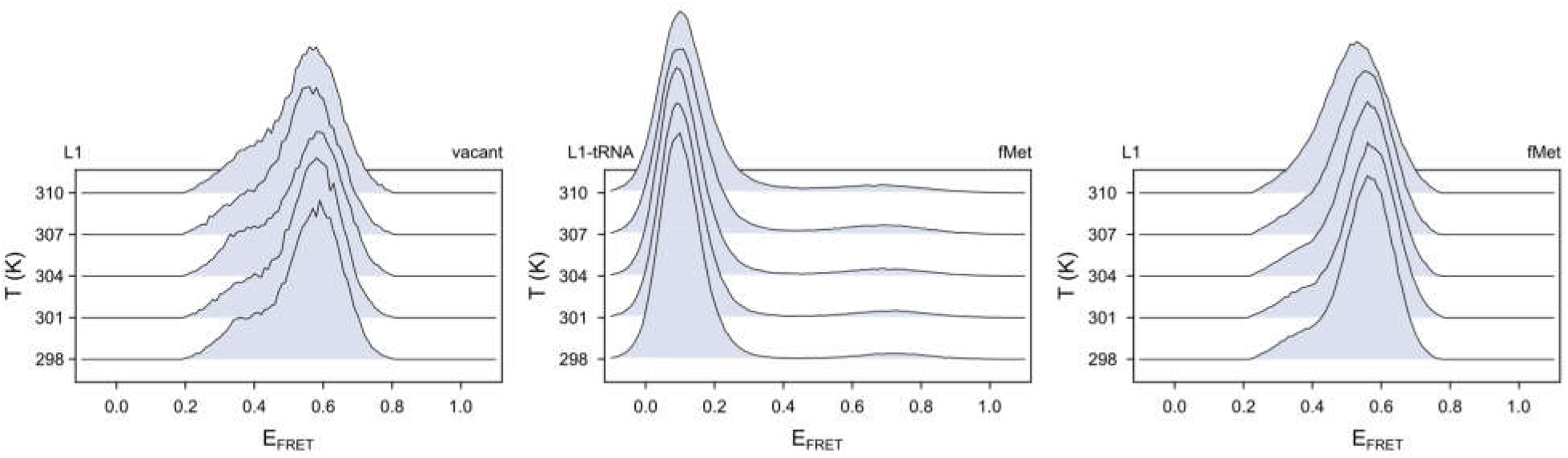
Histograms of smFRET signals for various RCs. Histograms of E_FRET_ values collected using smFRET_L1_ for RC^vacant^ (left), smFRET_L1-tRNA_ for RC^fMet^ (center), and smFRET_L1_ for RC^fMet^ (right) at five temperature points between 298 K and 310 K.

**Figure S4.**
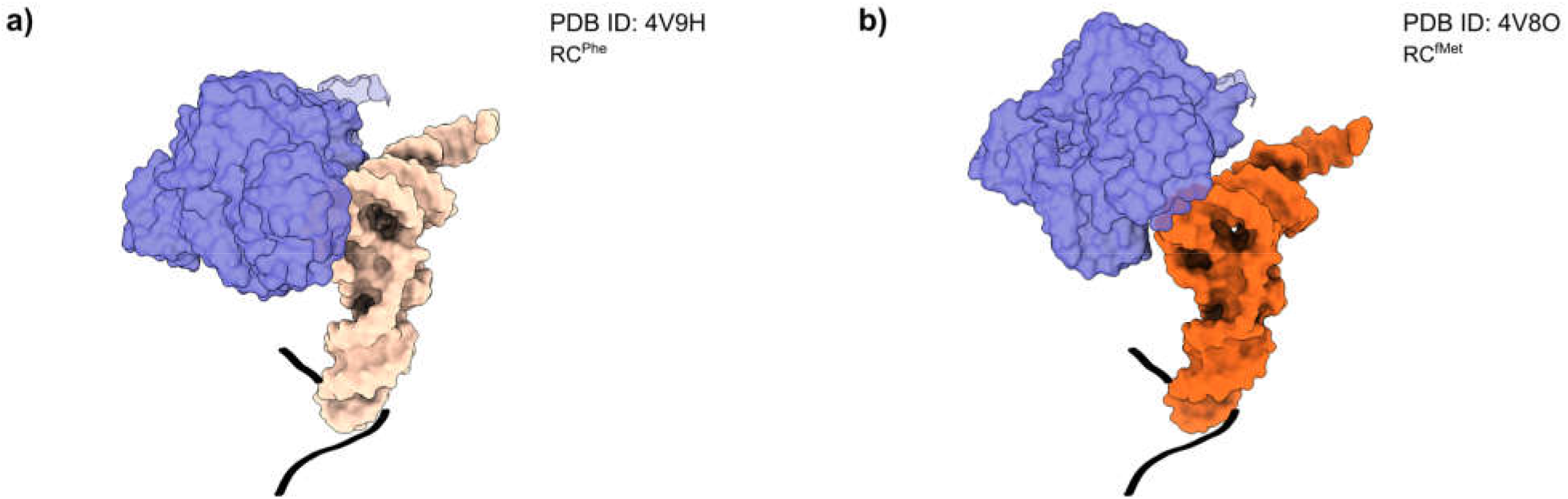
The conformations of the tRNA and L1 stalk in GS2 for RC^Phe^ and RC^fMet^. **(a)** The atomic structure of GS2 from an RC analogous to RC^Phe^ are compared to **(b)** the atomic structure of GS2 from an RC analogous to RC^fMet^. The two structures were positioned in the same orientation by aligning the anticodon stem loops of the tRNAs, were rendered in surface representations, and, for clarity, only the L1 stalk (the entire uL1 protein and nucleotides 2100-2200 of the 23S rRNA) (dark purple), tRNAs (tan for tRNA^Phe^ or orange for tRNA^fMet^), and path of the mRNA (black) are displayed.

**Figure S5.**
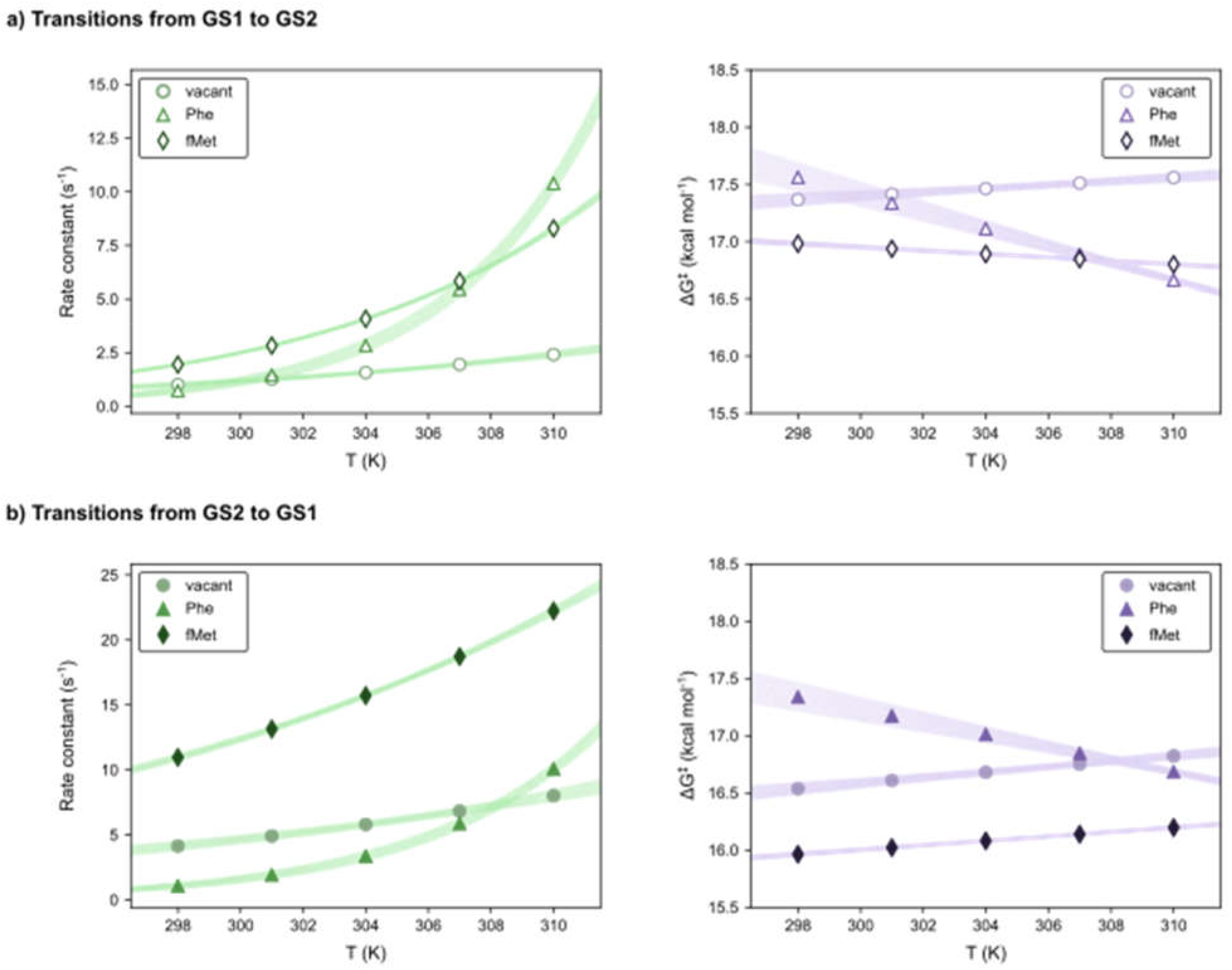
Transition rate constants and free energy barriers calculated from the activation parameters for all RCs. Transition rate constants (green) and the corresponding free energy barriers (*ΔG*^*‡*^) (purple) for the GS1→ GS2 (a) and GS2→ GS1 (b) transitions calculated at each of the five experimental temperature points from the global estimates of the activation enthalpies and entropies for RC^vacant^, RC^Phe^, and RC^fMet^ using smFRET_L1_. The shaded region represents the 95% confidence intervals for the rate constants for each RC, and the corresponding free energy barriers, calculated over the range of 297 K to 311 K using the corresponding posterior distributions for Δ*H*^*‡*^ and Δ*S*^*‡*^ (see Methods, and Table S1 for the number of individual molecules per dataset).

**Figure S6.**
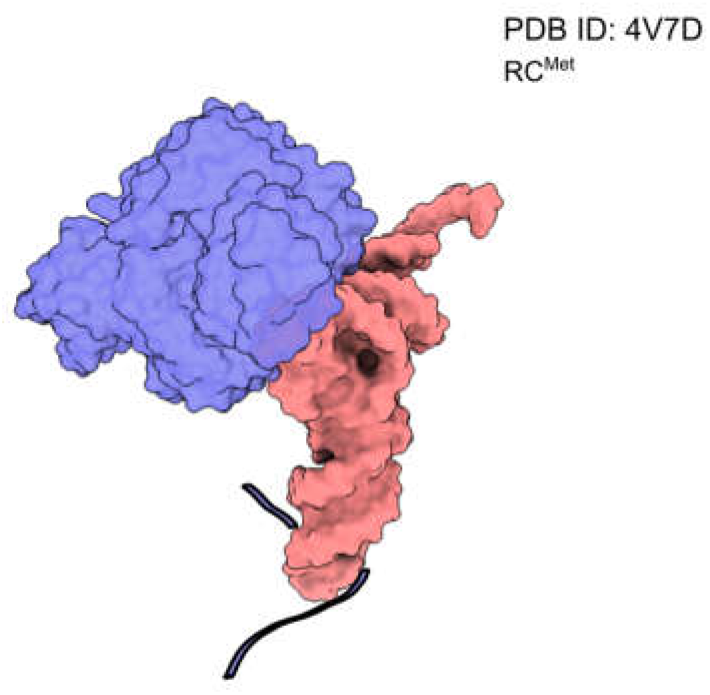
The conformation of the tRNA and L1 stalk in GS2 for RC^fMet^. Reconstructed atomic structure of an RC analogous to RC^Met^ in GS2. While the RC had been prepared using tRNA^Met^, the original structural model had been constructed using tRNA^Phe^. In this model, the tRNA has been replaced with tRNA^Met^, with a further round of rigid body refinement. The RC is positioned to the same orientation as the previous RCs (in Extended Data Figure 4) utilizing the alignment of the anticodon stem loops of the tRNAs. Similar to the previous structures, this structure was rendered in surface representation and, for clarity, only the L1 stalk (the entire uL1 protein and nucleotides 2100-2200 of the 23S rRNA) (dark purple), tRNA^Met^ (pink), and path of the mRNA (black) are displayed.

**Table S1.** The number of smFRET trajectories and total number of datapoints (in parentheses) used for analysis for each RC at each temperature.

|  | Trajectories (Datapoints) |  |  |  |  |
| --- | --- | --- | --- | --- | --- |
|  | 298 K | 301 K | 304 K | 307 K | 310 K |
| **RC<sup>vacant</sup>** | 340<br>(32321) | 296<br>(25829) | 363<br>(34887) | 369<br>(31874) | 366<br>(31958) |
| **RC<sup>Phe</sup> (L1)** | 590<br>(64281) | 536<br>(68000) | 543<br>(72474) | 542<br>(68879) | 387<br>(46057) |
| **RC<sup>Phe</sup> (L1-tRNA)** | 407<br>(111307) | 420<br>(128693) | 467<br>(151071) | 452<br>(128637) | 361<br>(105157) |
| **RC<sup>fMet</sup> (L1)** | 1769<br>(219410) | 1485<br>(183364) | 1540<br>(181060) | 1804<br>(191044) | 1739<br>(158692) |
| **RC<sup>fMet</sup> (L1-tRNA)** | 752<br>(294998) | 725<br>(334850) | 787<br>(287819) | 954<br>(334174) | 897<br>(306594) |

**Table S2.**
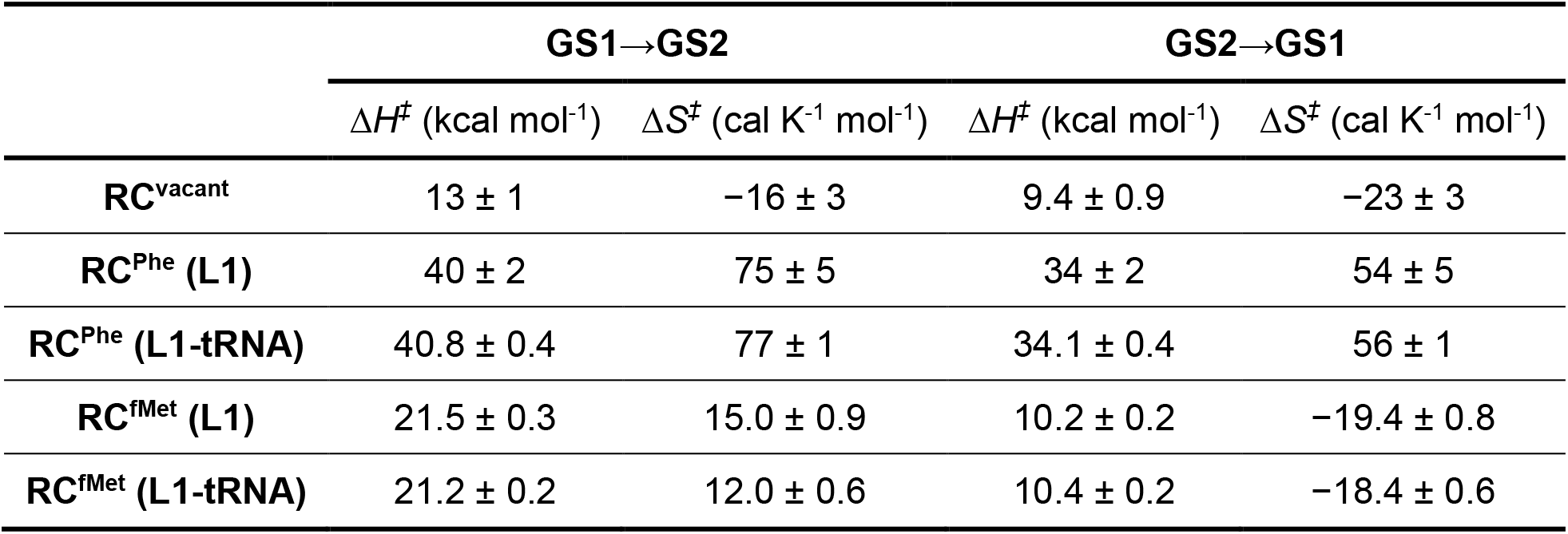
Activation parameter values for all RCs. The absolute values of activation parameters for each RC. Errors are standard deviations of the marginalized posterior probability distributions of each parameter (see Methods, and Table S1 for the number of individual molecules per dataset).

**Table S3.** The prior probability distributions used for the parameters during the global BIASD analysis. *ϵ*_*GS1*_ and *ϵ*_*GS2*_ are the mean E_FRET_s for GS1 and GS2 respectively; *σ* is the noise for both states; *ΔH*_^*‡*^(GS1→ GS2) and_ *ΔH*_^*‡*^(GS2→ GS1)_ are the enthalpic components of the free energy barriers for the GS1→ GS2 and GS2→ GS1 transitions respectively; and *ΔS*^*‡*^_(GS1→ GS2) and_ *ΔS*_^*‡*^(GS2→ GS1)_ are the corresponding entropic components. *μ* and *τ* are the mean and standard deviations of the Gaussian distribution, while *a* and *b* are the minimum and maximum values of the uniform distribution.

| Parameter | Dataset | Distribution | Distribution Parameters |
| --- | --- | --- | --- |
| $\epsilon_{GS1}$ | smFRET<sub>L1-tRNA</sub> | Gaussian | $\mu = 0.15, \tau = 0.1$ |
| | smFRET<sub>L1</sub> (RC<sup>Phe</sup>) | Gaussian | $\mu = 0.55, \tau = 0.1$ |
| | smFRET<sub>L1</sub> (RC<sup>vacant</sup> and RC<sup>fMet</sup>) | Gaussian | $\mu = 0.5840, \tau = 0.0001$ |
| $\epsilon_{GS2}$ | smFRET<sub>L1-tRNA</sub> | Gaussian | $\mu = 0.85, \tau = 0.1$ |
| | smFRET<sub>L1</sub> (RC<sup>Phe</sup>) | Gaussian | $\mu = 0.35, \tau = 0.1$ |
| | smFRET<sub>L1</sub> (RC<sup>vacant</sup> and RC<sup>fMet</sup>) | Gaussian | $\mu = 0.3545, \tau = 0.0001$ |
| $\sigma$ | All | Uniform | $a = 0.0001, b = 0.15$ |
| $\Delta H^\ddagger_{(\text{GS1} \rightarrow \text{GS2})}$ | All | Uniform | $a = -5 \times 10^5, b = 5 \times 10^5$ |
| $\Delta S^\ddagger_{(\text{GS1} \rightarrow \text{GS2})}$ | All | Uniform | $a = -1000, b = 1000$ |
| $\Delta H^\ddagger_{(\text{GS2} \rightarrow \text{GS1})}$ | All | Uniform | $a = -5 \times 10^5, b = 5 \times 10^5$ |
| $\Delta S^\ddagger_{(\text{GS2} \rightarrow \text{GS1})}$ | All | Uniform | $a = -1000, b = 1000$ |

## Notes

https://github.com/KorakRay/biasd-tst

### Competing Interest Statement

The authors have declared no competing interest.

### Summary of Updates

The Introduction section has been revised to include three additional references. The Abstract has been revised with minor changes. The manuscript has been updated with minor changes in formatting.

